# StableLift: Optimized Germline and Somatic Variant Detection Across Genome Builds

**DOI:** 10.1101/2024.10.31.621401

**Authors:** Nicholas K. Wang, Nicholas Wiltsie, Helena K. Winata, Sorel Fitz-Gibbon, Alfredo E. Gonzalez, Nicole Zeltser, Raag Agrawal, Jieun Oh, Jaron Arbet, Yash Patel, Takafumi N. Yamaguchi, Paul C. Boutros

**Author notes:** Corresponding author: Center for Health Sciences, 10833 Le Conte Avenue, Los Angeles, CA, United States of America 90095.

## Abstract

Reference genomes are foundational to modern genomics. Our growing understanding of genome structure leads to continual improvements in reference genomes and new genome “builds” with incompatible coordinate systems. We quantified the impact of genome build on germline and somatic variant calling by analyzing tumour-normal whole-genome pairs against the two most widely used human genome builds. The average individual had a build-discordance of 3.8% for germline SNPs, 8.6% for germline SVs, 25.9% for somatic SNVs and 49.6% for somatic SVs. Build-discordant variants are not simply false-positives: 47% were verified by targeted resequencing. Build-discordant variants were associated with specific genomic and technical features in variant- and algorithm-specific patterns. We leveraged these patterns to create StableLift, an algorithm that predicts cross-build stability with AUROCs of 0.934 ± 0.029. These results call for significant caution in cross-build analyses and for use of StableLift as a computationally efficient solution to mitigate inter-build artifacts.

## Main

Since initial assembly of the human genome in 2001^1,2^, thousands of errors have been corrected, polymorphic regions have been defined and the diversity of included individuals has expanded^3–5^. These advances have led to a series of updated human reference genome “builds”, each with incompatible coordinate numbering. While new builds are more accurate representations, their adoption can be slow in both research and clinical settings^6^.

One key factor slowing adoption of new genome builds is computational cost: re-aligning sequencing data requires local storage of raw reads and investment of substantial compute time. To avoid these time and financial costs, tools have been created to convert or “liftover” genomic coordinates between builds^7,8^. Despite widespread use, coordinate conversion using these tools was designed for larger intervals and can introduce artifacts when applied to individual variant calls^9–17^. It remains unclear whether and what biases are introduced by coordinate conversion, especially in the context of structural and somatic variant detection.

To fill this gap, we compared DNA whole genome sequencing (WGS) alignment and variant detection on the two most widely used reference genomes: GRCh37 and GRCh38 (**Figure 1a**). Fifty human tumour-normal WGS pairs were analyzed on both builds using identical tools and software versions *via* standardized Nextflow pipelines (**Supplementary Figure 1a; Supplementary Table 1**)^18–20^. Variants detected from sequencing data aligned to GRCh37 were converted to GRCh38 coordinates using BCFtools/liftover^21^ with UCSC chain files^22,23^. Converted GRCh37 variants were compared to variants detected from sequencing data directly aligned to GRCh38. We evaluated four variant classes: germline single nucleotide polymorphisms (gSNPs, including indels), germline structural variants (gSVs), somatic single nucleotide variants (sSNVs, including indels) and somatic structural variants (sSVs).

**Figure 1:**
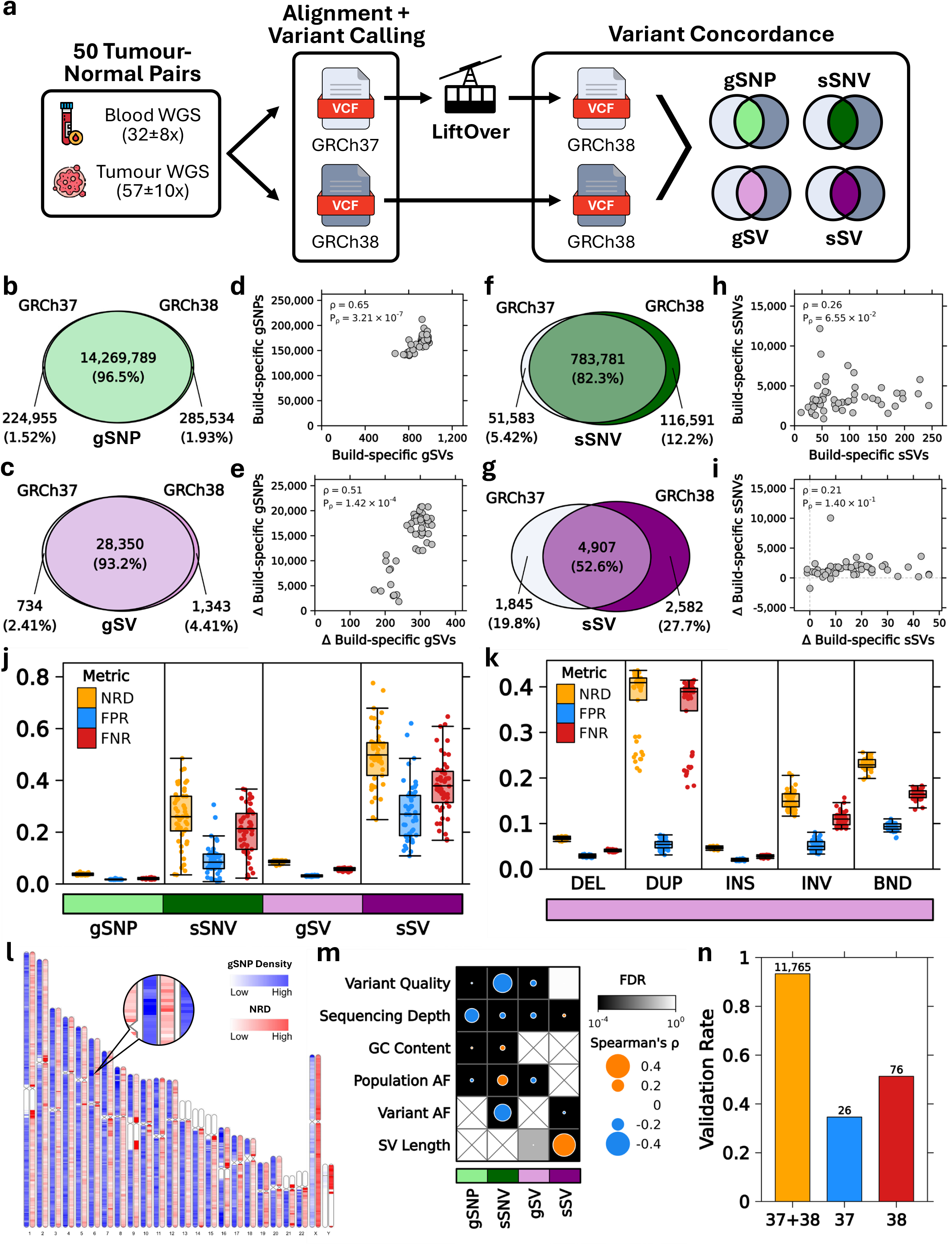
Overview of differences between GRCh37 and GRCh38 variant calls. **a)** Experimental design for matched comparison of germline and somatic variants in a representative cancer genomics workflow. **b-c)** Cohort level overlap of converted GRCh37 *vs*. GRCh38 germline variants (gSNP, gSV). **d)** Number of build-specific germline variants per sample. **e)** Difference in per sample germline variant counts found in GRCh38 relative to GRCh37. **f-g)** Cohort level overlap of converted GRCh37 *vs*. GRCh38 somatic variants (sSNV, sSV). **h)** Number of build-specific somatic variants per sample. **i)** Difference in per sample somatic variant counts found in GRCh38 relative to GRCh37. **j-k)** Variant discordance per sample stratified by variant type and gSV subtype. (NRD = non-reference discordance, FPR = false positive rate, FNR = false negative rate; DEL = deletion, DUP = duplication, INS = insertion, INV = inversion, BND = breakend/translocation) **l**) Distribution of gSNP density and NRD across the genome. **m)** Correlation between continuous covariates and NRD per variant type. Spearman’s correlation indicated by dot size and color; statistical significance with false discovery rate correction indicated by background shading. **n)** Validation rate of build-concordant, GRCh37-specific, and GRCh38-specific sSNVs by targeted deep-sequencing.

Most germline SNPs and structural variants identified were shared between the two builds (>93%; **Figure 1b-c**). Nevertheless, we detected 166,704 ± 14,829 build-specific gSNPs and 908 ± 73 build-specific gSVs per individual (mean ± standard deviation; **Figure 1d**). Alignment to GRCh38 led to identification of more gSNPs and gSVs (**Figure 1e**). By contrast, somatic variant detection was dramatically more variable: only 82% of sSNVs and 53% of sSVs were identified in both builds (**Figure 1f-g**). This led to 3,611 ± 2,025 build-specific sSNVs and 93 ± 61 build-specific sSVs (**Figure 1h**), with more somatic variants identified when aligning to GRCh38 (**Figure 1i**).

To better characterize build-specific calls, we calculated three complementary metrics of genotype concordance. First, we assessed non-reference discordance (NRD), which is the fraction of all non-reference genotypes that disagree between builds. Next, we considered direct variant calling on GRCh38 as ground truth and calculated false positive rate (FPR) and false negative rate (FNR). Consistent with variant detection numbers, all three metrics of genotype concordance were substantially better for germline than somatic variants: 3.8 ± 0.0% NRD for gSNPs and 8.6 ± 0.1% for gSVs *vs*. 25.9 ± 11.0% for sSNVs and 49.6 ± 11.2% for sSVs (per individual mean ± standard deviation; **Figure 1j**). The high FNR of somatic variant detection on GRCh37 (20.4 ± 9.5% sSNVs, 38.1 ± 11.0% sSVs; **Figure 1j**) suggests that the many published studies aligning to GRCh37 may systematically underestimate somatic mutation burden (or alternatively those aligning to GRCh38 may overestimate it).

To understand whether these discordances are randomly distributed, we first evaluated different classes of gSVs. Deletions and insertions were less discordant between builds than duplications, inversion and translocations (**Figure 1k**). The high FNR of duplications (35.2 ± 7.7%) suggested increased sensitivity in GRCh38 potentially due to improved resolution of duplicated or homologous regions. This led us to investigate whether discordance in germline SNPs also varied spatially across the genome. Consistent with the gSV results, we observed significant heterogeneity in build-specific differences within and across chromosomes (**Figure 1l**). For example, a one Mbp region of 6p21.3 in the HLA region contained 16,784 gSNPs with mean 8.5% NRD, while a neighboring one Mbp region had 8,626 gSNPs with mean 1.2% NRD.

A wide range of other features are associated with discordance across builds (**Figure 1m**; **Supplementary Figure 1-7**). As an example, discordant sSNVs were more likely to have lower quality scores but higher GC content (**Figure 1m**; **Supplementary Figure 2a**,**c**). Discordant sSNVs also exhibited a non-monotonic association with coverage: both atypically-low and atypically-high coverage was associated with increased discordance, possibly due to erroneous mapping to homologous or repetitive regions (**Supplementary Figure 2b**). sSNVs with higher somatic allele frequencies tended to be less discordant, while variants seen at higher allele frequencies in TOPMed^24^ were more likely to be discordant (**Figure 1m**; **Supplementary Figure 2d-e**). Discordance rates varied significantly across chromosomes (mean NRD ranging from 6.3% on chromosome 13 to 47.8% on chromosome Y; **Supplementary Figure 6a**) and trinucleotide contexts (mean NRD ranging from 4.7% to 17.3%; **Supplementary Figure 6d**). sSNVs in satellite repeat regions were particularly discordant (mean 59.8% NRD; **Supplementary Figure 6e**), supportive of repetitive regions as a major source of discordance.

One natural explanation of these results is that almost all build-discordant genetic variation results from false-positive predictions from variant-detection algorithms. To quantify this, we exploited targeted deep-sequencing validation (mean 653x coverage) on sSNV calls from five tumour-normal, whole genome pairs (**Supplementary Table 2**)^25^. Build-concordant variants had a validation rate of 93.3% (**Figure 1n**). Nevertheless, 34.6% of GRCh37-specific variants and 51.3% of GRCh38-specific variants were validated by targeted deep-sequencing. This is a clear enrichment of false-positives relative to build-concordant variants, but demonstrates that build-specific variants are a balance of false-positives and false-negative predictions. As a result, simply using the latest genome build is insufficient: one third of variants detected on GRCh37 but not in GRCh38 are false-negatives.

To quantify the cross-build stability of any individual variant, we created a machine-learning approach called StableLift. By leveraging features associated with build-discordance (**Supplementary Figures 1-7**), StableLift estimates the likelihood (“Stability Score”) that a given variant will be consistently represented across two genome builds (**Figure 2a**). We trained StableLift with variants detected from the same fifty tumour-normal WGS pairs using six variant callers spanning all four variant-types: HaplotypeCaller^26^, MuTect2^27^, Strelka2^28^, SomaticSniper^29^, MuSE2^30^ and DELLY2^31^. We validated StableLift in 10 tumour-normal whole genomes^32^ (**Supplementary Table 3**) and 60 tumour-normal exomes^32^ (**Supplementary Table 4**) for area under the receiver operating characteristic curve (AUROC) and selected a default operating point to maximize F_1_-score in the whole genome validation set.

**Figure 2:**
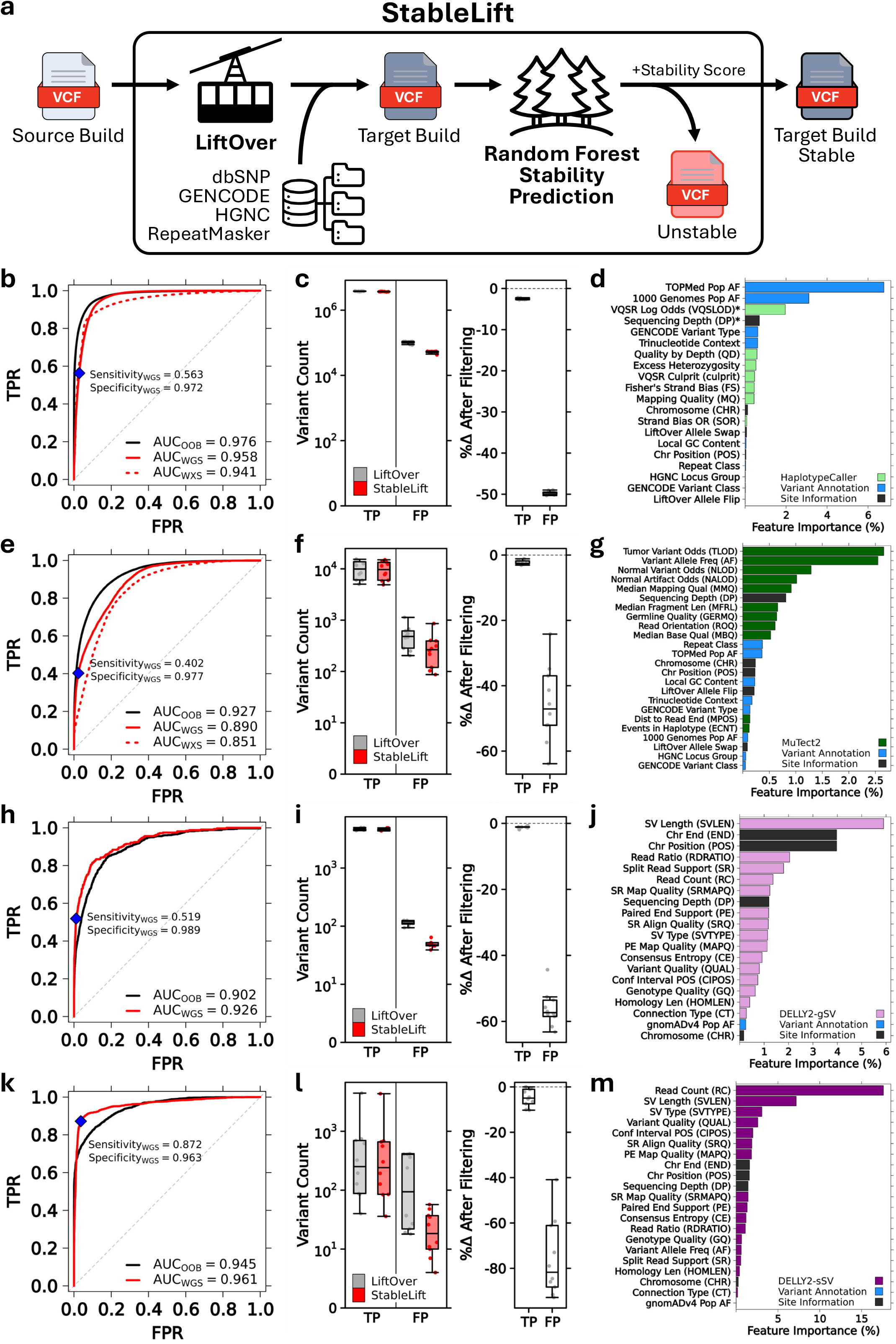
Machine-learning approach to predicting variant stability across genome builds. **a)** Overview of StableLift as a multi-purpose genomics utility performing LiftOver coordinate conversion, variant annotation, and cross-build stability prediction. **b)** Random forest model performance for gSNPs (HaplotypeCaller) shown as ROC curves and AUC measures for out-of-bag whole genome training (OOB, solid black), whole genome validation (WGS, solid red), and whole exome validation (WXS, dashed red) sets. Default operating point maximizing F_1_-score highlighted (blue diamond) with corresponding sensitivity and specificity in the whole genome validation set. **c)** Comparison of concordant (TP) and discordant (FP) gSNP counts before and after default StableLift filtering. **d)** Random forest feature importance colored by caller-specific metrics, variant annotations and site information. Normalized features indicated by *. **e-g)** Same as b-d for sSNVs (MuTect2). **h-j)** Same as b-d for gSVs (DELLY2). **k-m)** Same as b-d for sSVs (DELLY2).

StableLift robustly identified build-discordant gSNP calls, with validation AUROCs of 0.958 for WGS and 0.941 for exome sequencing (**Figure 2b; Supplementary Figure 8a-c**). At the F_1_-maximizing operating point, 49.7 ± 0.5% of discordant gSNPs in WGS validation were discarded, corresponding to 51,181 ± 4,884 discordant variants removed per individual (**Figure 2c**). A variety of features contributed to the accuracy of these predictions, most notably TOPMed^24^ population allele frequency (**Figure 2d**) driven by elevated discordance of variants with allele frequencies near zero (rare variants/singletons) or one (reference artifacts; **Supplementary Figure 1e**).

StableLift similarly identified build-discordant sSNVs, with validation AUROCs of 0.890 for WGS and 0.851 for exome sequencing (MuTect2; **Figure 2e; Supplementary Figure 8d-f**) and a 45.7 ± 11.7% reduction of discordant calls (−209 ± 56 discordant sSNVs; **Figure 2f**). sSNV stability prediction was driven by a wide range of predictor features (**Figure 2g**). Models fit to three other sSNV callers achieved similar performance: AUROC_WGS_ = 0.932 for Strelka2, AUROC_WGS_ = 0.964 for SomaticSniper and AUROC_WGS_ = 0.905 for MuSE2 (**Supplementary Figure 9a-i, Supplementary Figure 10**). Different sSNV calling algorithms had similar but not identical patterns of feature importance, highlighting the interaction between genomic features and variant detection algorithms (**Supplementary Figure 9j**).

To understand how predicted variant stability relates to variant validation status, we ran StableLift on the previously described five whole genome pairs with targeted deep-sequencing validation (**Supplementary Figure 11a**). sSNVs predicted to be “Stable” were 1.3-9.6x more likely to validate than those predicted to be “Unstable” (**Supplementary Figure 11b-c**). Similarly, the Stability Score distribution was higher for validated *vs*. unvalidated variants (**Supplementary Figure 11d-g**).

Finally, we applied StableLift to structural variant calls made by DELLY2^31^. Despite only 28,350 concordant cases and 734 discordant cases of gSV training data (**Figure 1c**), StableLift again accurately identified discordant calls, with a validation AUROC of 0.926 (**Figure 2h**) and a 56.2 ± 5.3% reduction of discordant calls (−63 ± 10 discordant gSVs; **Figure 2i**). Length of variant was the most important single feature, with a range of predictive features differing from those driving the accuracy of the gSNP and sSNV models (**Figure 2j**). Accuracy in DELLY2 sSVs was equally high, achieving a validation AUROC of 0.961 (**Figure 2k**) and removing 81.7% of discordant sSVs (−171 ± 170 discordant sSVs; **Figure 2l**). Only 4,907 concordant and 1,845 discordant training cases were needed for this model, and its accuracy was driven by read count and SV length (**Figure 2m**).

This work calls for significant caution in cross-build analyses. GRCh37 remains in routine use and while re-alignment to GRCh38 is preferable, this is computationally expensive. In many cases realignment may not be possible: raw data or software pipelines may no longer be available, particularly for older technologies. Similarly, variant databases created with GRCh37 coordinates can introduce challenges in annotating newer GRCh38-derived results. StableLift can create models to convert between any two genome builds. While our results focused on converting GRCh37 results to GRCh38, we provide models of similar accuracy for the inverse conversion of GRCh38 to GRCh37 (**Supplementary Figure 12-16**).

StableLift provides an attractive approach to mitigate bias in many cases, but the build-sensitivity of somatic and structural variant calling warrants increased attention from algorithm developers. Some biases appear to be systematic, and while GRCh38 calls are generally more accurate, we identified apparent false-negatives with both genome builds. As genetic analyses gradually transition from linear reference genomes to graph-based pangenomes^33–38^, quantifying build-specific variation and efficiently minimizing error rates in cross-build conversion will become increasingly important.

## Online Methods

### Analysis cohort

To assess LiftOver concordance in a representative cancer genomics workflow, we chose to evaluate a cohort of 50 patients spanning eight cancer types from the International Cancer Genome Consortium (ICGC PRAD-CA)^18^ and the Pan-Cancer Analysis of Whole Genomes (PCAWG) Consortium^25^ (**Supplementary Table 1**). All patients had paired tumour-normal whole-genome sequencing with germline and somatic coverage of 32±8x and 57±10x, respectively.

### Alignment and variant calling

Sequencing reads were aligned to the GRCh37 (hs37d5) and GRCh38 (hg38) reference builds using BWA-MEM2 (v2.2.1)^39^ in paired-end, alt-aware mode followed by GATK’s ‘MarkDuplicatesSpark’ (v4.2.4.1)^26^ (**Supplementary Figure 1a**). Indel realignment and base quality score recalibration were performed using GATK’s ‘IndelRealigner’ (v3.7.0), ‘BaseRecalibrator’ (v4.2.4.1), and ‘ApplyBQSR’ (v4.2.4.1)^26^. Germline SNPs were called using GATK’s ‘HaplotypeCaller’ (v4.2.4.1) in GVCF mode followed by variant quality score recalibration using ‘VariantRecalibrator’ (v4.2.4.1) and ‘ApplyVQSR’ (v4.2.4.1) and joint genotyping across all normal samples using ‘GenotypeGVCFs’ (v4.2.4.1)^26^. Somatic SNVs were called using MuTect2 (v4.2.4.1)^27^ in tumour-normal mode with default parameters. Germline and somatic SVs were called using DELLY2 (v1.2.6)^31^ with default parameters and a more stringent minimum paired-end mapping quality threshold of 20. Germline SVs were regenotyped using the output of ‘delly merge’ and filtered with ‘delly filter -f germline’ (v1.2.6)^31^.

All alignment and variant calling operations were run on a Slurm high-performance computing cluster using Nextflow (v23.04.2) pipelines^19,20,40^ to ensure reproducibility and compatibility across computing environments. The GRCh37 and GRCh38 analysis pipelines used identical parameters except for the reference genome input and associated resource files.

### LiftOver coordinate conversion

GRCh37 SNV calls were converted to GRCh38 coordinates using the BCFtools/liftover plugin (v1.20)^21^ with UCSC chain files^22,23^. For SVs, a custom R script was used to convert variants by breakpoint (CHROM, POS, END for DEL, DUP, INS, INV variants; CHROM, POS, END, CHR2, POS2 for BND variants) using the UCSC chain files along with the rtracklayer (v1.62.0)^41^ and GenomicRanges (v1.54.1)^42^ R packages.

### Variant concordance

SNV concordance was evaluated at the cohort level using ‘vcf-compare’ from VCFtools (v0.1.16)^43^ and at the sample level using ‘SnpSift concordance’ (v5.2.0)^44^. Per variant SNV concordance was quantified using ‘bcftools stats --verbose’ (v1.20)^45^. SV concordance was evaluated using ‘SV Concordance’ (v4.4.0.0) from GATK.

To accurately assess the practical impacts of LiftOver operations on variant calling, performance metrics need to be carefully chosen^46^. Metrics including true negative counts should be used with caution. In the case of SNVs, the number of sites matching the reference far outnumber variant sites and can lead to inflated estimates of accuracy. Furthermore, standard SNV calling pipelines typically only report sites which differ from the reference sequence. Outside of targeted re-genotyping, the absence of a variant cannot be assumed to be a reference match as the missing call could be attributed to a lack of coverage or insufficient evidence. This issue is even more pronounced with structural variants.

We utilized the following three metrics to i) characterize the concordance and error profiles of LiftOver operations and ii) provide guidance for when and where these errors are the most relevant. True positive (TP), false positive (FP), true negative (TN) and false negative (FN) calls are computed for converted GRCh37 variant calls relative to GRCh38.

Non-reference discordance (NRD) measures the overall disagreement between the two variant sets and is equivalent to overall accuracy with true negatives excluded from the denominator:

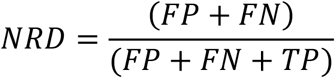

False positive rate (FPR) represents the fraction of variants identified in GRCh37, but not in GRCh38:

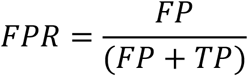

False negative rate (FNR) represents the fraction of variants identified in GRCh38, but not in GRCh37:

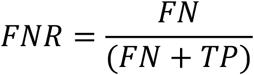

### Variant annotation

For SNVs, dbSNP (build 151)^47^, GENCODE (v34)^48^, and HGNC (Nov302017)^49^ annotations were added using GATK’s ‘Funcotator’ (v4.6.0.0)^26^ with pre-packaged data source v1.7.20200521s. Trinucleotide context was determined using ‘bedtools getfasta’ (v2.31.0)^50^. RepeatMasker (v3.0.1)^51^ intervals were obtained from the UCSC Table Browser^22^ and intersected with variant calls using ‘bedtools intersect’ (v2.31.0)^50^. SVs were intersected with the gnomAD-SV (v4)^52^ database (FILTER == “PASS”) using a custom R script and annotated with population allele frequency.

### Targeted sequencing validation

Additional targeted deep-sequencing data from five patients in the analysis cohort^25,53^ (653x mean coverage; **Supplementary Table 2**) was used to validate a subset of sSNV calls. sSNVs identified in the whole genome data within targeted validation regions were considered validated if they were also identified in the targeted deep-sequencing data (**Supplementary Figure 11a**).

### Random forest stability prediction

Using the variant calls from our analysis cohort and their corresponding NRD labels, we trained a random forest model to predict variant concordance for each of six variant callers – HaplotypeCaller (v4.2.4.1)^26^, MuTect2 (v4.2.4.1)^27^, Strelka2 (v2.9.10)^28^, SomaticSniper (v1.0.5.0)^29^, MuSE2 (v2.0.4)^30^, DELLY2 (v1.2.6)^31^ – across four variant types (gSNP, sSNV, gSV, sSV; **Supplementary Figure 1a**). Variants were dichotomized based on a 20% NRD threshold and a probability forest (’num.trees’ = 500 for gSNPs and 1,000 for sSNVs, gSVs, sSVs) was trained using the ranger (v0.16.0)^54^ R package to predict concordant *vs*. discordant variants. Variants failing LiftOver coordinate conversion were excluded. The model outputs a “Stability Score” for each variant indicating the fraction of trees predicting concordant status.

### Feature selection and hyperparameter optimization

The set of features considered for each model included all variant fields provided by each variant caller, along with external annotations and site information. Feature inclusion and normalization were determined by optimizing for AUROC in the validation sets for each respective model. Hyperparameters were tuned using a grid search over ‘mtry’ and ‘min.node.size’.

### Model validation datasets

For gSNPs and sSNVs, 10 sarcoma tumour-normal whole genome pairs (**Supplementary Table 3**) and 60 sarcoma tumour-normal exome pairs (**Supplementary Table 4**) from The Cancer Genome Atlas (TCGA-SARC)^32^ were used as validation sets to demonstrate generalizability across sequencing methods (whole genome *vs*. exome) and cancer types (sarcoma not represented in the training set). Raw sequencing data was downloaded and reprocessed with the same pipelines used for the comparative analysis. For gSVs and sSVs, only the 10 whole genome pairs were used for validation as exome data provides insufficient coverage for comprehensive SV calling.

Five whole genomes from the targeted sequencing validation cohort^25,53^ were used to evaluate StableLift predictions against an independent truth set of validated *vs*. unvalidated sSNVs (**Supplementary Table 2; Supplementary Figure 11a**).

### StableLift

We incorporated these pre-trained and validated models into a standardized workflow accepting either GRCh37 or GRCh38 input VCFs from six variant callers (HaplotypeCaller, MuTect2, Strelka2, SomaticSniper, MuSE2, DELLY2) spanning four variant types (gSNP, sSNV, gSV, sSV). Input variants are converted and annotated as described above and output with a predicted “Stability Score” for filtering based on user-specified thresholds. Performance in the TCGA-SARC whole genome validation set is included with each model to define the default F_1_-maximizing operating point and allow for custom filtering based on pre-calibrated sensitivity and specificity estimates.

### Data visualization

Figures were generated in R (v4.3.3) using the lattice (v0.22-6), latticeExtra (v0.6-30), BPG (v7.1.0)^55^, VennDiagram (v1.7.3)^56^, and RIdeogram (v0.2.2)^57^ packages.

## Data availability

Somatic VCFs, resource files for variant annotation, and pre-trained random forest models for GRCh37→GRCh38 and GRCh38→GRCh37 conversions are available on GitHub as release attachments (https://github.com/uclahs-cds/pipeline-StableLift/releases). The tumour-normal whole genome pairs used for analysis and training StableLift can be accessed through the European Genome-Phenome Archive (https://ega-archive.org/studies/EGAS00001000900) and the Bionimbus Protected Data Cloud (https://icgc.bionimbus.org/). TCGA-SARC exome and whole genome datasets used for validation can be accessed from the GDC Data Portal (portal.gdc.cancer.gov/projects/TCGA-SARC).

## Code availability

StableLift is available on GitHub (https://github.com/uclahs-cds/pipeline-StableLift) as a Nextflow pipeline featuring LiftOver coordinate conversion, variant annotation with external databases and prediction of cross-build variant stability. Nextflow pipelines for alignment and variant calling are on GitHub (https://github.com/uclahs-cds/metapipeline-DNA) and described elsewhere^20^.

## Acknowledgments

The authors thank all members of the Boutros lab and the Office of Health Informatics and Analytics (OHIA) at UCLA. The comparative analysis and training of StableLift were based upon data generated by the International Cancer Genome Consortium (ICGC) and the Pan-Cancer Analysis of Whole Genomes (PCAWG) Consortium. Validation of StableLift was based upon data generated by The Cancer Genome Atlas (TCGA) Research Network and the Pan-Cancer Analysis of Whole Genomes (PCAWG) Consortium.

## Author Contributions

**Conceptualization:** NKW, PCB

**Data Curation:** NKW, SF, AEG, RA, JO, YP, TNY

**Formal Analysis:** NKW, NZ

**Funding Acquisition:** PCB

**Methodology:** NKW, HKW, JA, PCB

**Software:** NKW, NW, YP

**Supervision:** PCB

**Writing – Original Draft:** NKW, PCB

**Writing – Review & Editing:** NKW, NW, HKW, SF, AEG, NZ, RA, JO, JA, YP, TNY, PCB

