## Supplementary Figures for "StableLift: Optimized Germline and Somatic Variant Detection Across Genome Builds"

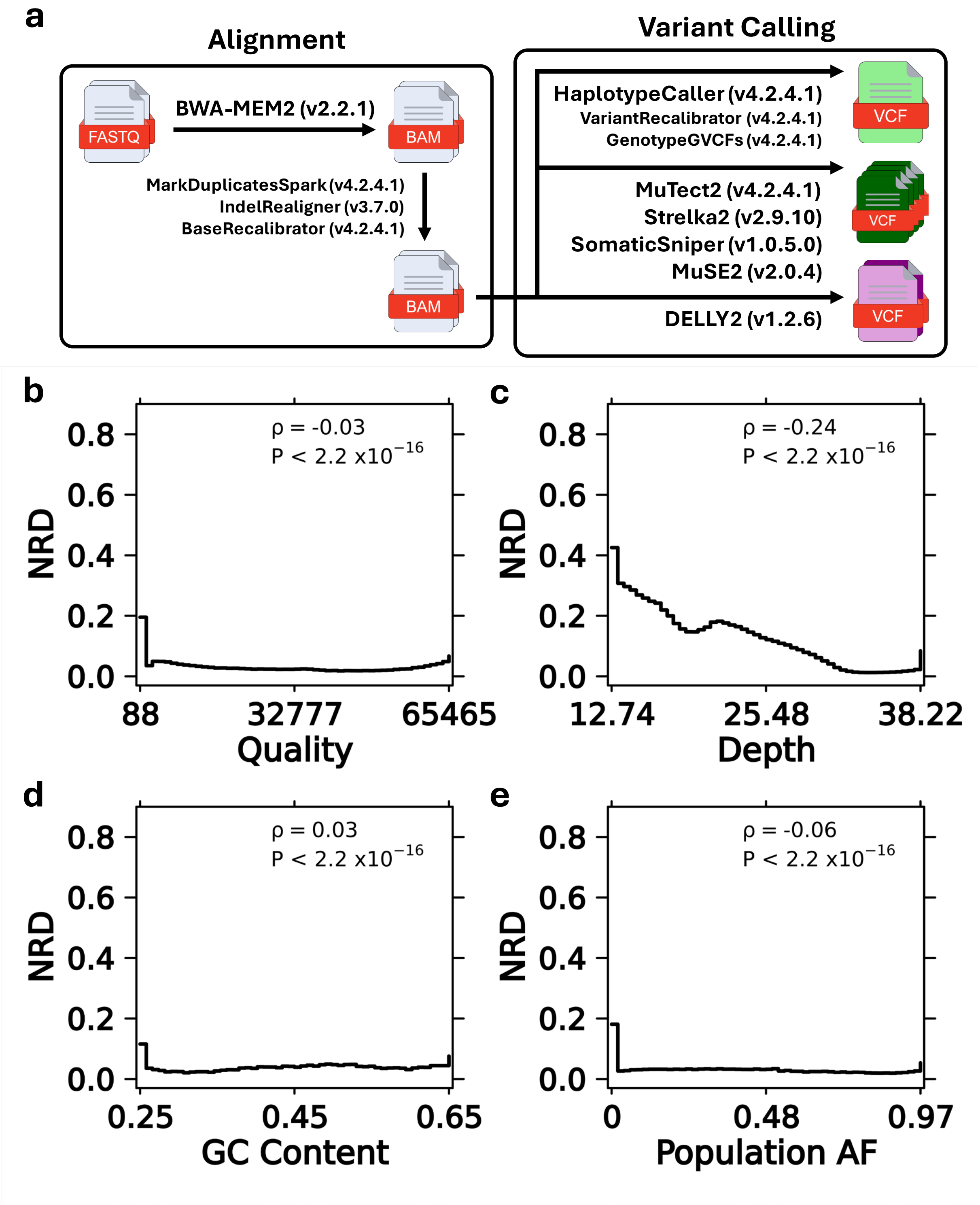


**Supplementary Figure 1. a)** Overview of alignment and variant calling pipelines for gSNPs (HaplotypeCaller), sSNVs (MuTect2, Strelka2, SomaticSniper, MuSE2), gSVs (DELLY2), and sSVs (DELLY2). **b-e)** Distribution of gSNP (HaplotypeCaller) non-reference discordance (NRD) across continuous covariates. Spearman correlation coefficients shown with corresponding p-values.

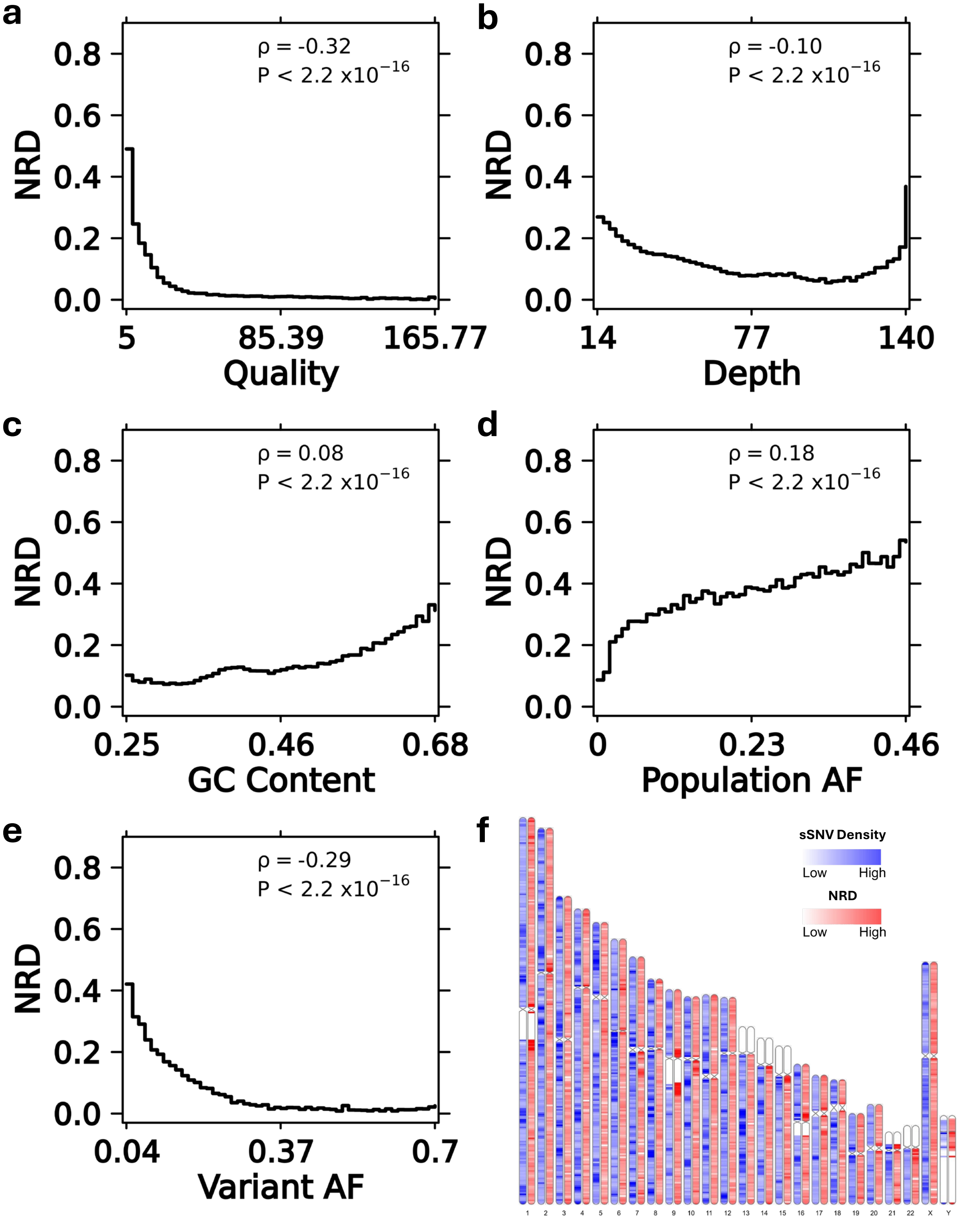


**Supplementary Figure 2. a-e)** Distribution of sSNV (MuTect2) non-reference discordance (NRD) across continuous covariates. Spearman correlation coefficients shown with corresponding p-values. **f)** Distribution of sSNV (MuTect2) density and NRD rates across the genome.

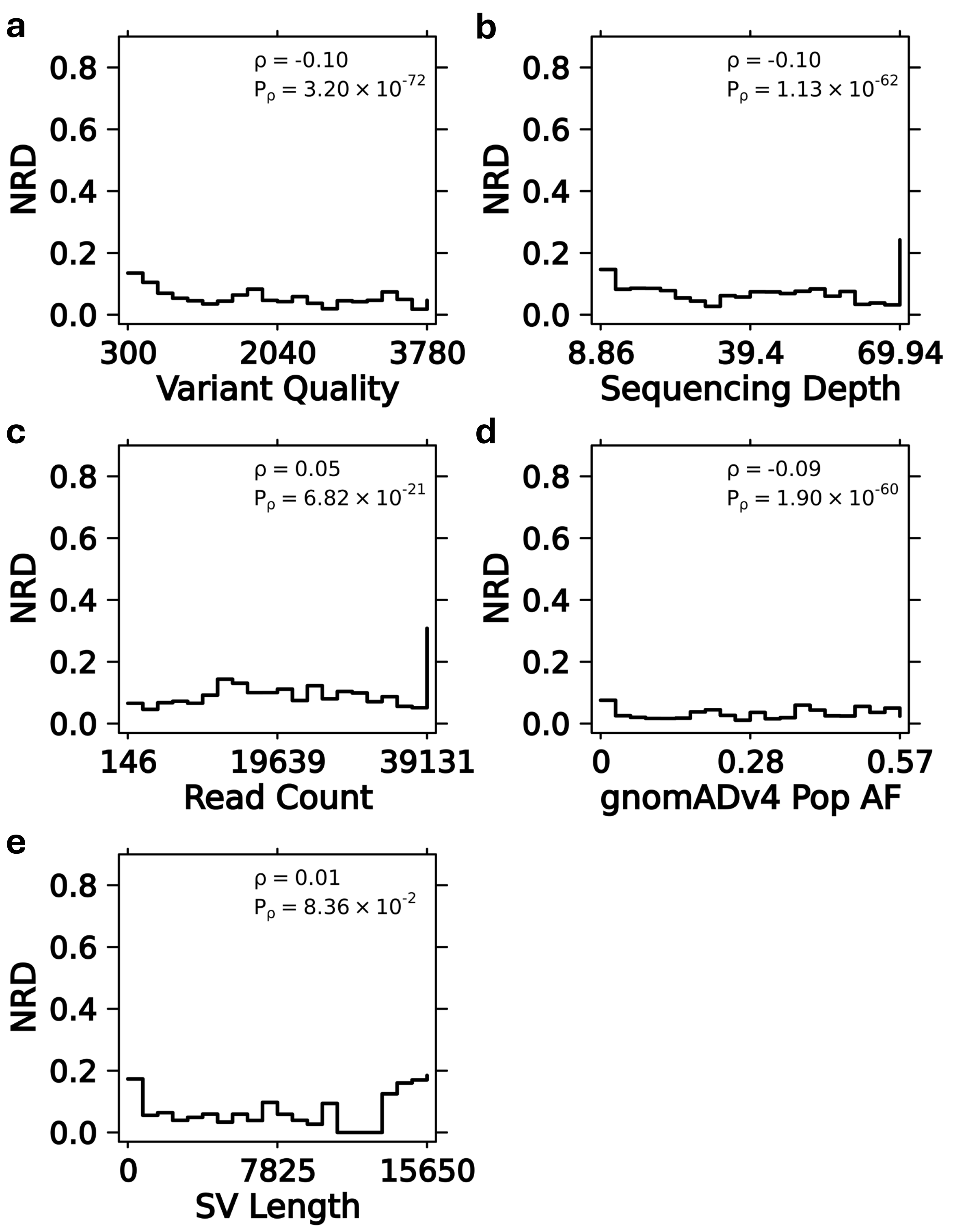


**Supplementary Figure 3. a-e)** Distribution of gSV (DELLY2) non-reference discordance (NRD) across continuous covariates. Spearman correlation coefficients shown with corresponding p-values.

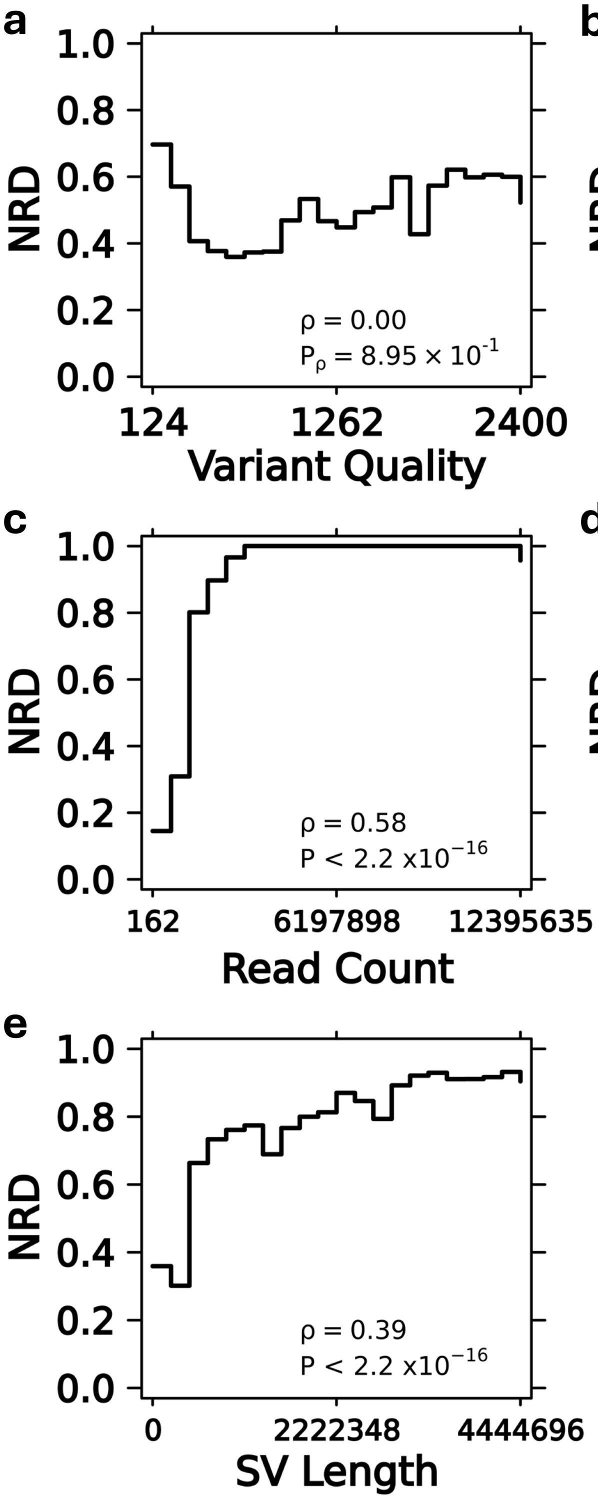


**Supplementary Figure 4. a-e)** Distribution of sSV (DELLY2) non-reference discordance (NRD) across continuous covariates. Spearman correlation coefficients shown with corresponding p-values.

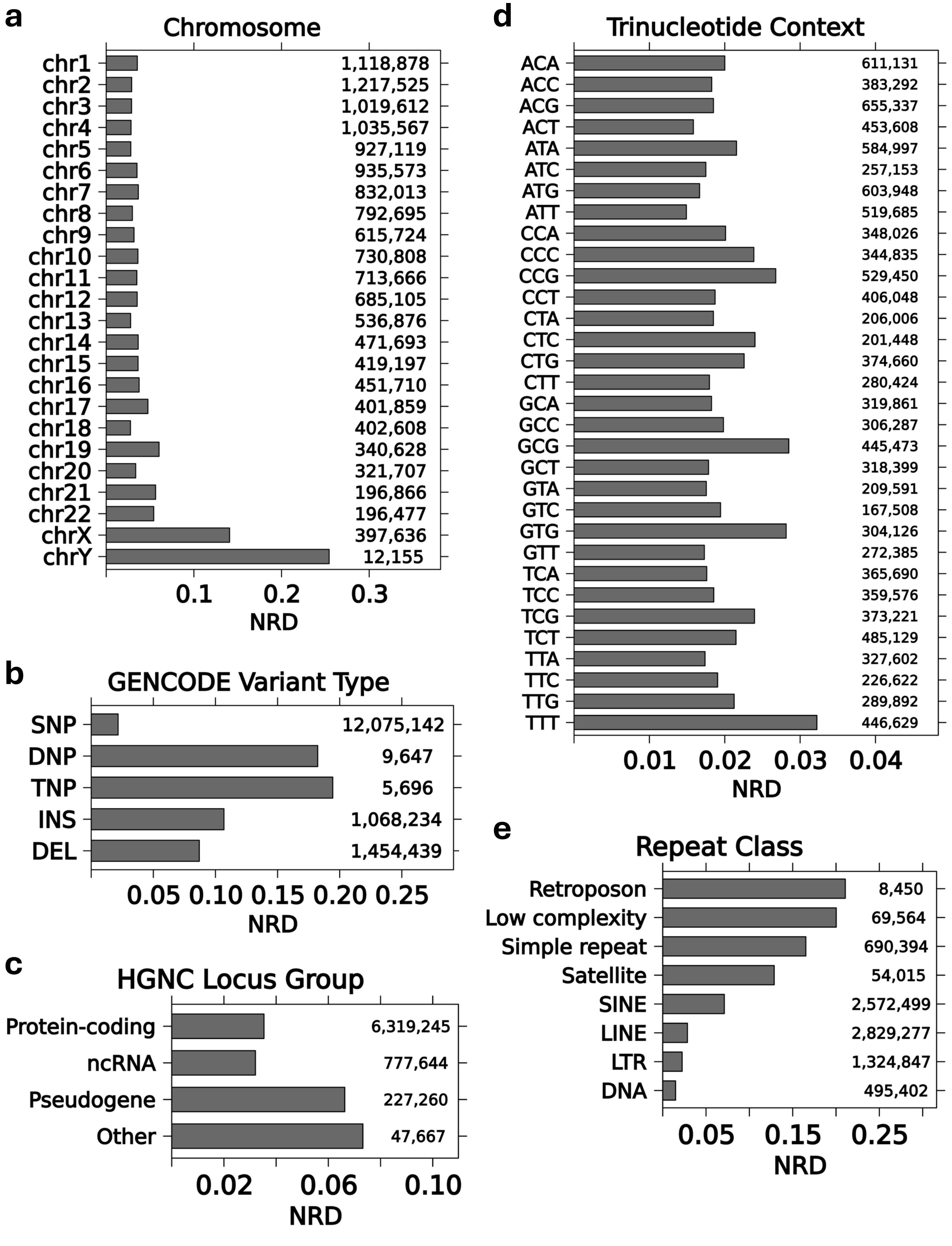


**Supplementary Figure 5. a-e)** Mean non-reference discordance (NRD) of gSNPs (HaplotypeCaller) across categorical covariates. Numeric label indicates number of variants per category.

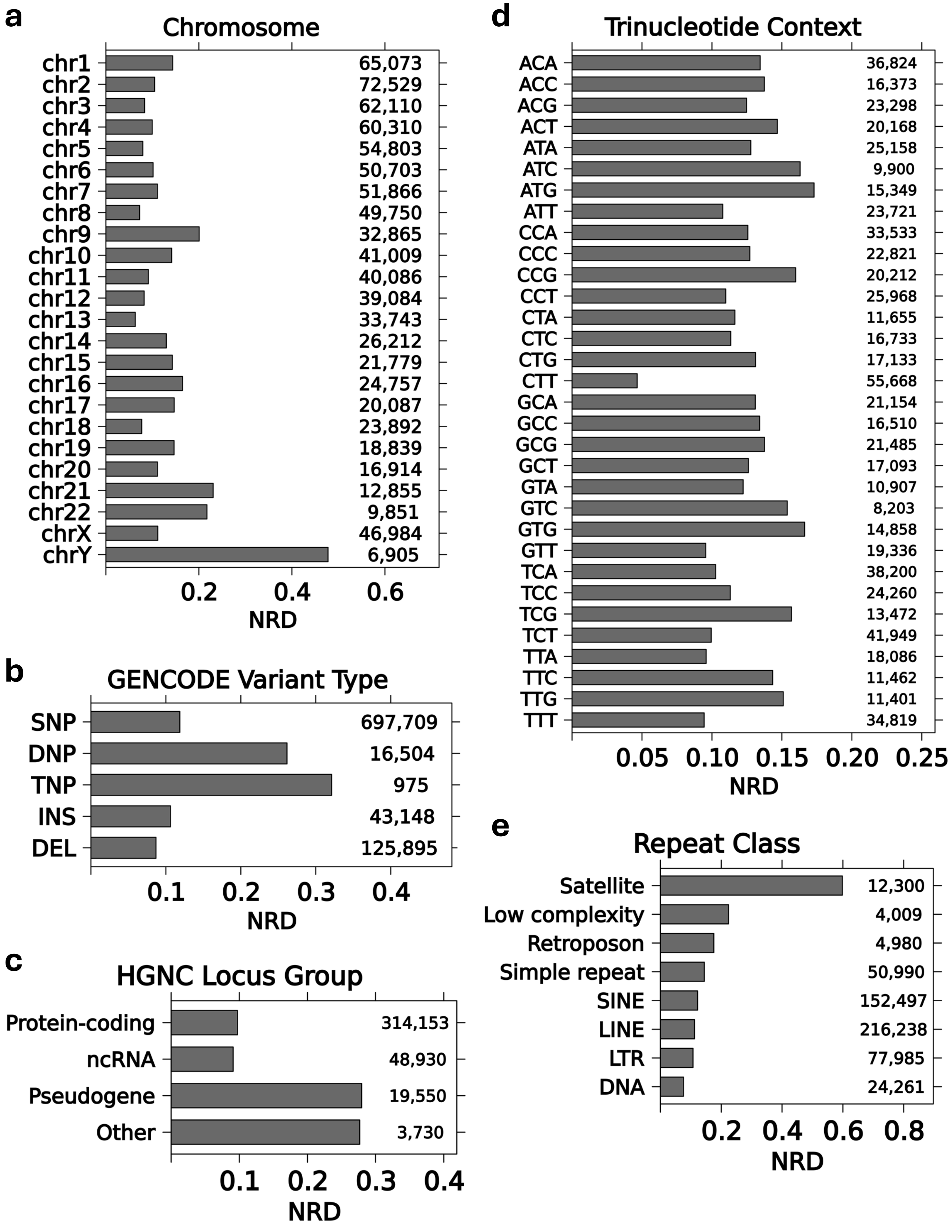


**Supplementary Figure 6. a-e)** Mean non-reference discordance (NRD) of sSNVs (MuTect2) across categorical covariates. Numeric label indicates number of variants per category.

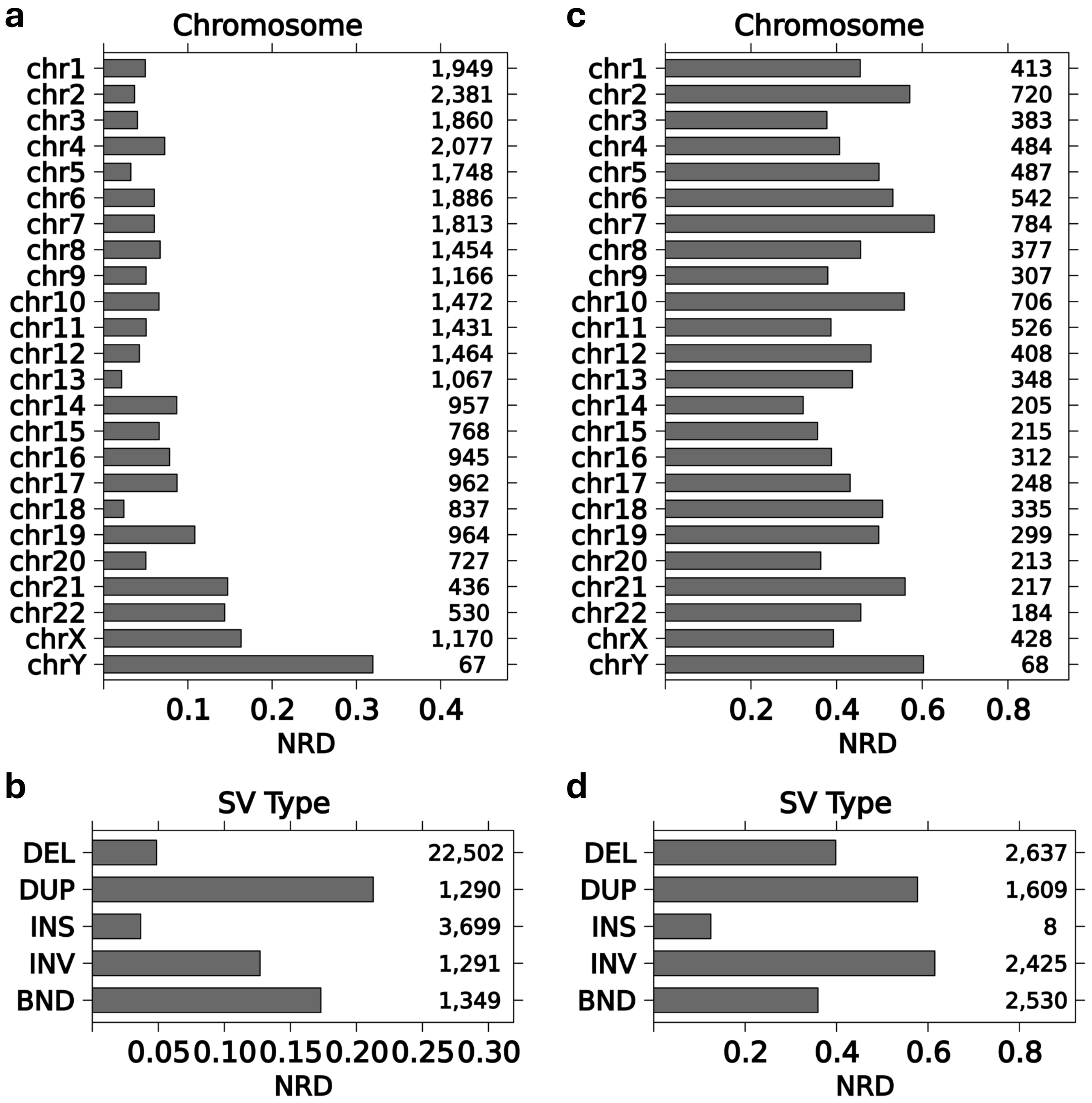


**Supplementary Figure 7. a-b)** Mean non-reference discordance (NRD) of gSVs (DELLY2) across categorical covariates. **c-d)** Mean non-reference discordance (NRD) of sSVs (DELLY2) across categorical covariates. Numeric label indicates number of variants per category.

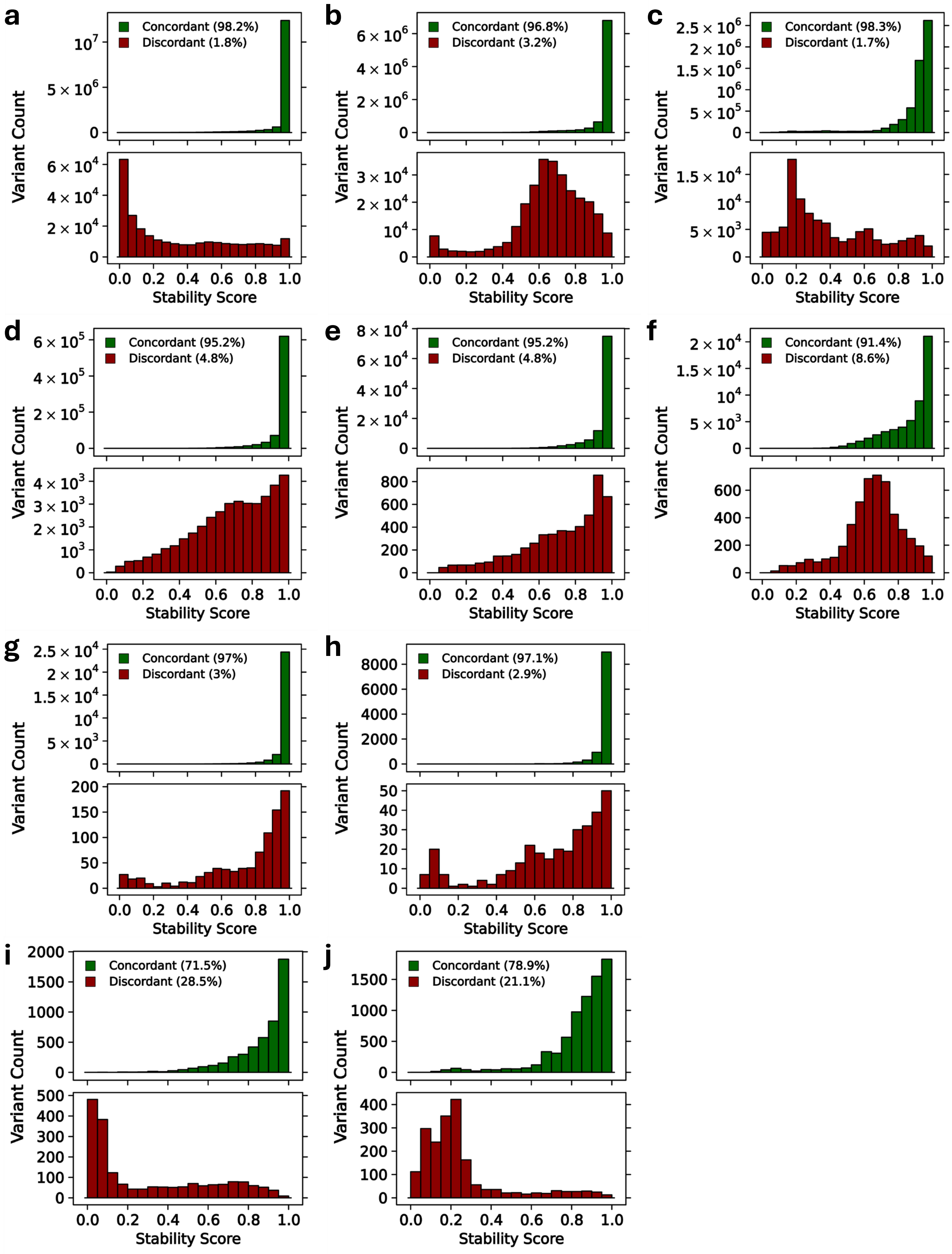


**Supplementary Figure 8.** GRCh37→GRCh38: Stability Score distribution of concordant *vs*. discordant variants (relative class proportions in parentheses) per variant caller. **a-c)** Concordant *vs*. discordant HaplotypeCaller gSNPs in whole genome training set (n=50), whole genome validation set (n=10), and exome validation set (n=60), respectively. **d-f)** Same as a-c for concordant *vs*. discordant MuTect2 sSNVs. **g-h)** Same as a-b for concordant *vs*. discordant DELLY2 gSVs. **i-j)** Same as a-b for concordant *vs*. discordant DELLY2 sSVs.

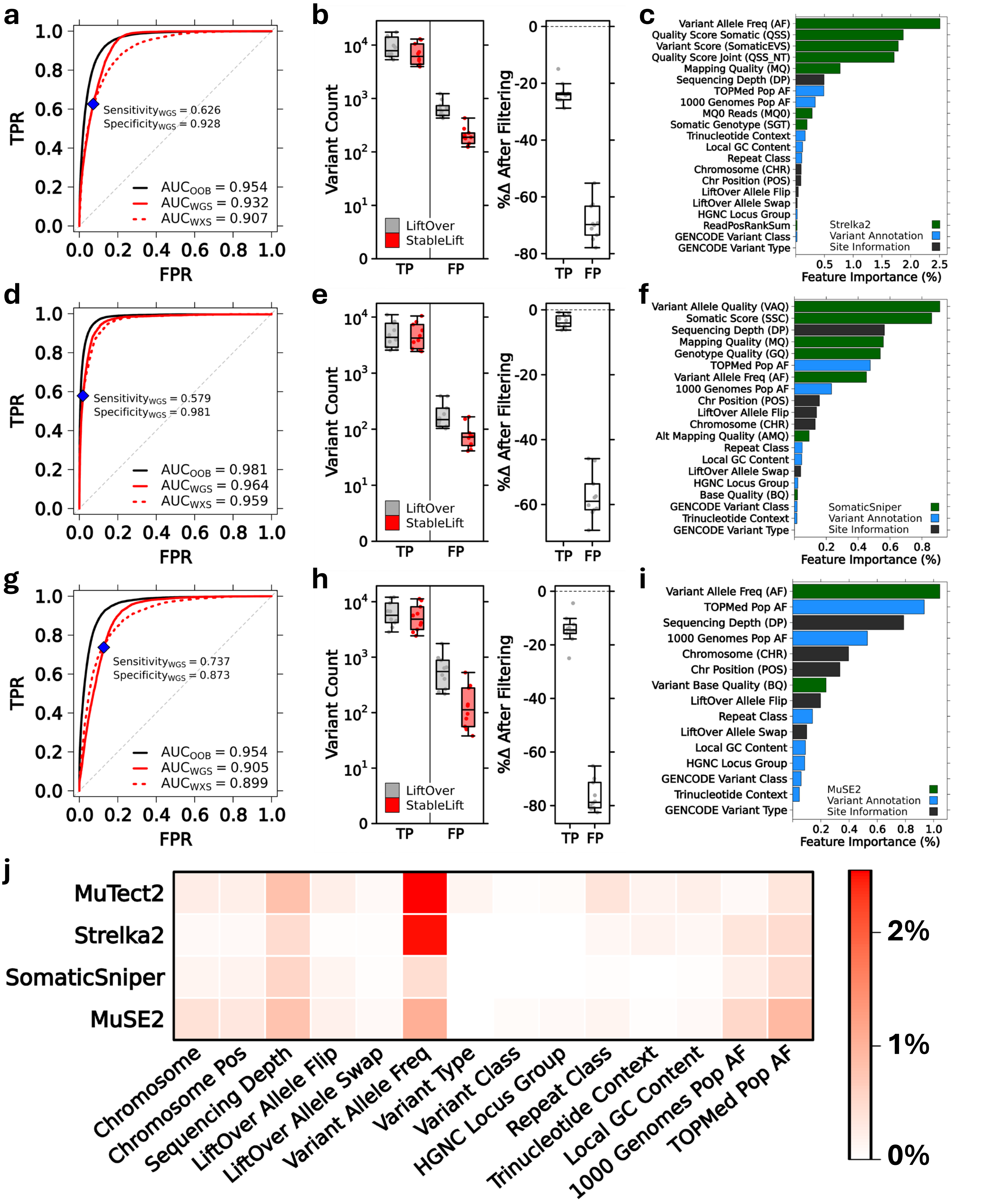


**Supplementary Figure 9.** GRCh37→GRCh38 StableLift model metrics. **a)** Random forest model performance for Strelka2 sSNVs shown as ROC curves and AUC measures for out-of-bag whole genome training (OOB, solid black), whole genome validation (WGS, solid red), and whole exome validation (WXS, dashed red) sets. Default operating point maximizing F1-score highlighted (blue) with corresponding sensitivity and specificity in the whole genome validation set. **b)** Comparison of concordant (TP) and discordant (FP) sSNV counts before and after default StableLift filtering. **c)** Random forest feature importance colored by caller-specific metrics, variant annotations, and site information. **d-f)** Same as a-c for SomaticSniper sSNVs. **g-i)** Same as a-c for MuSE2 sSNVs. **j)** Comparison of feature importance scores for features shared across the four sSNV callers.

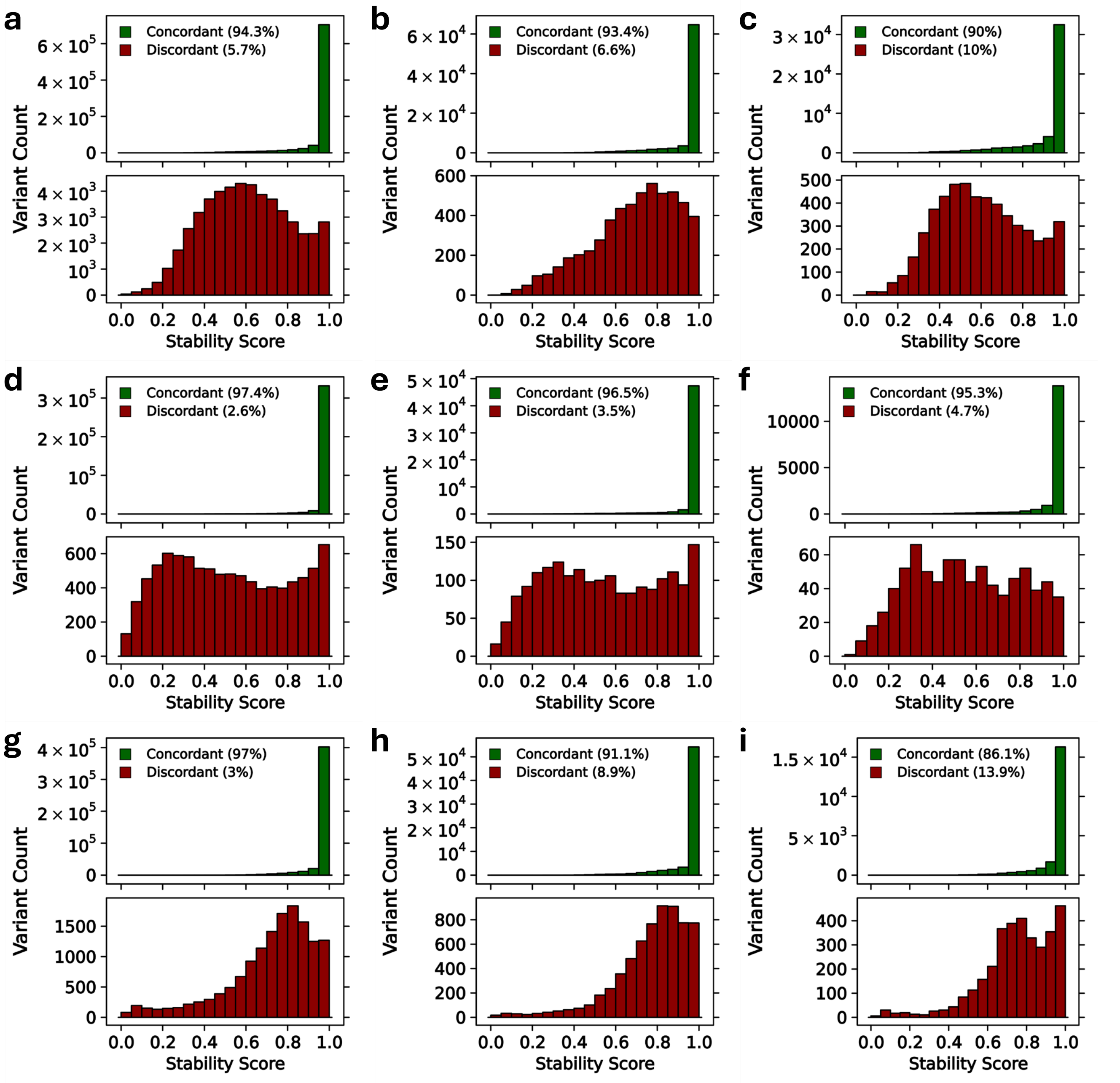


**Supplementary Figure 10.** GRCh37→GRCh38: Stability Score distribution of concordant *vs.* discordant variants (relative class proportions in parentheses) per variant caller. **a-c)** Concordant *vs.* discordant Strelka2 sSNVs in whole genome training set (n=50), whole genome validation set (n=10), and exome validation set (n=60), respectively. **d-f)** Same as a-c for concordant *vs.* discordant SomaticSniper sSNVs. **g-i)** Same as a-c for concordant *vs.* discordant MuSE2 sSNVs.

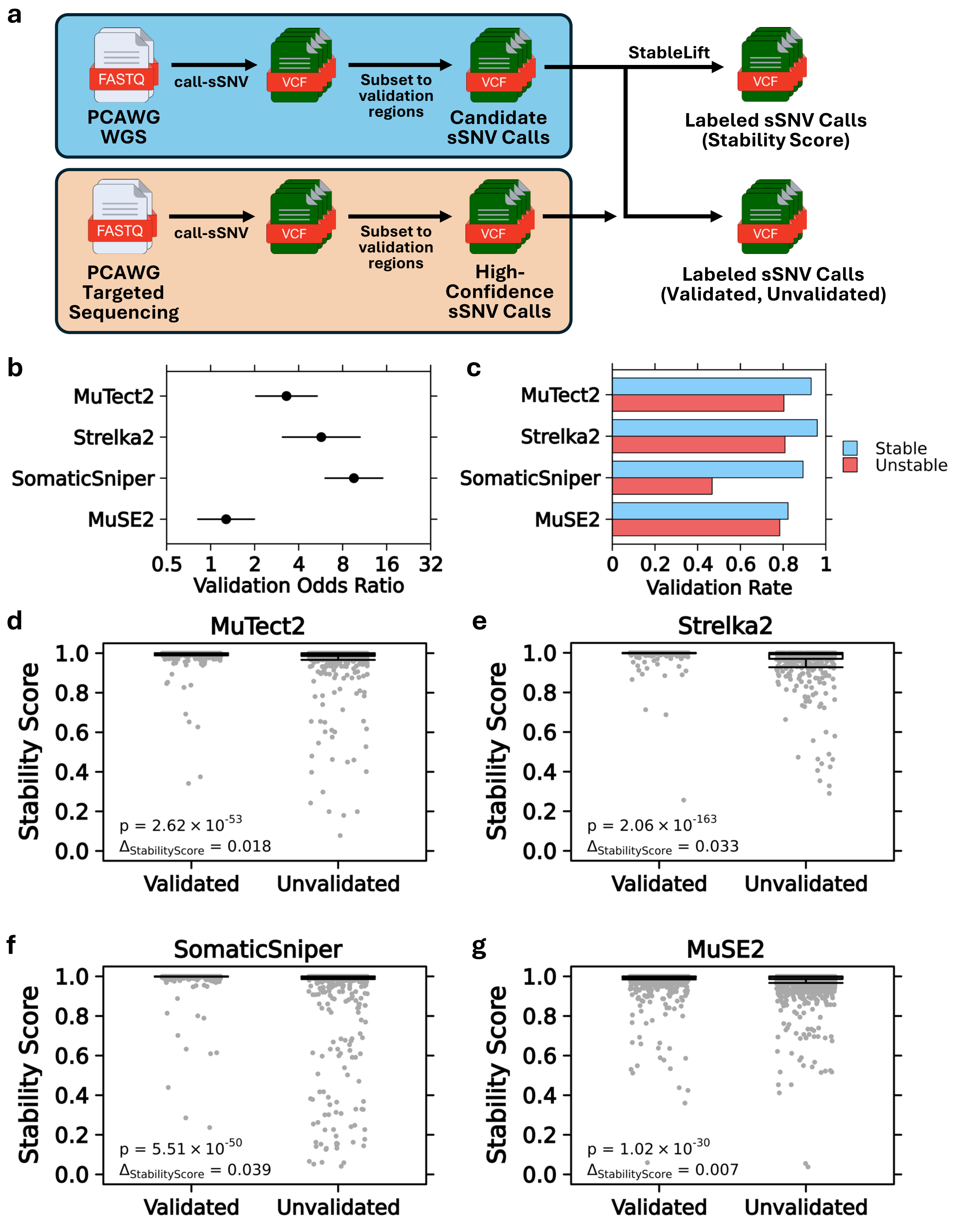


**Supplementary Figure 11.** GRCh37→GRCh38: Validation of StableLift on sSNV calls from five whole genome pairs with deep targeted resequencing. **a)** Experimental design comparing candidate sSNV calls from whole-genome sequencing with high-confidence sSNV calls from targeted deep-sequencing. **b)** Validation odds ratio with 95% confidence interval of predicted "Stable" *vs*. "Unstable" variants for each sSNV caller. Validation status determined by concordance with variant calls from targeted deep-sequencing. **c)** Validation rate of predicted "Stable" *vs*. "Unstable" variants for each sSNV caller. **d-g)** Distribution of Stability Scores for validated *vs*. unvalidated sSNVs. Stability Scores per variant caller with Wilcoxon test p-value and difference in mean Stability Score shown. Strip plot downsampled to normalize group sizes for visualization.

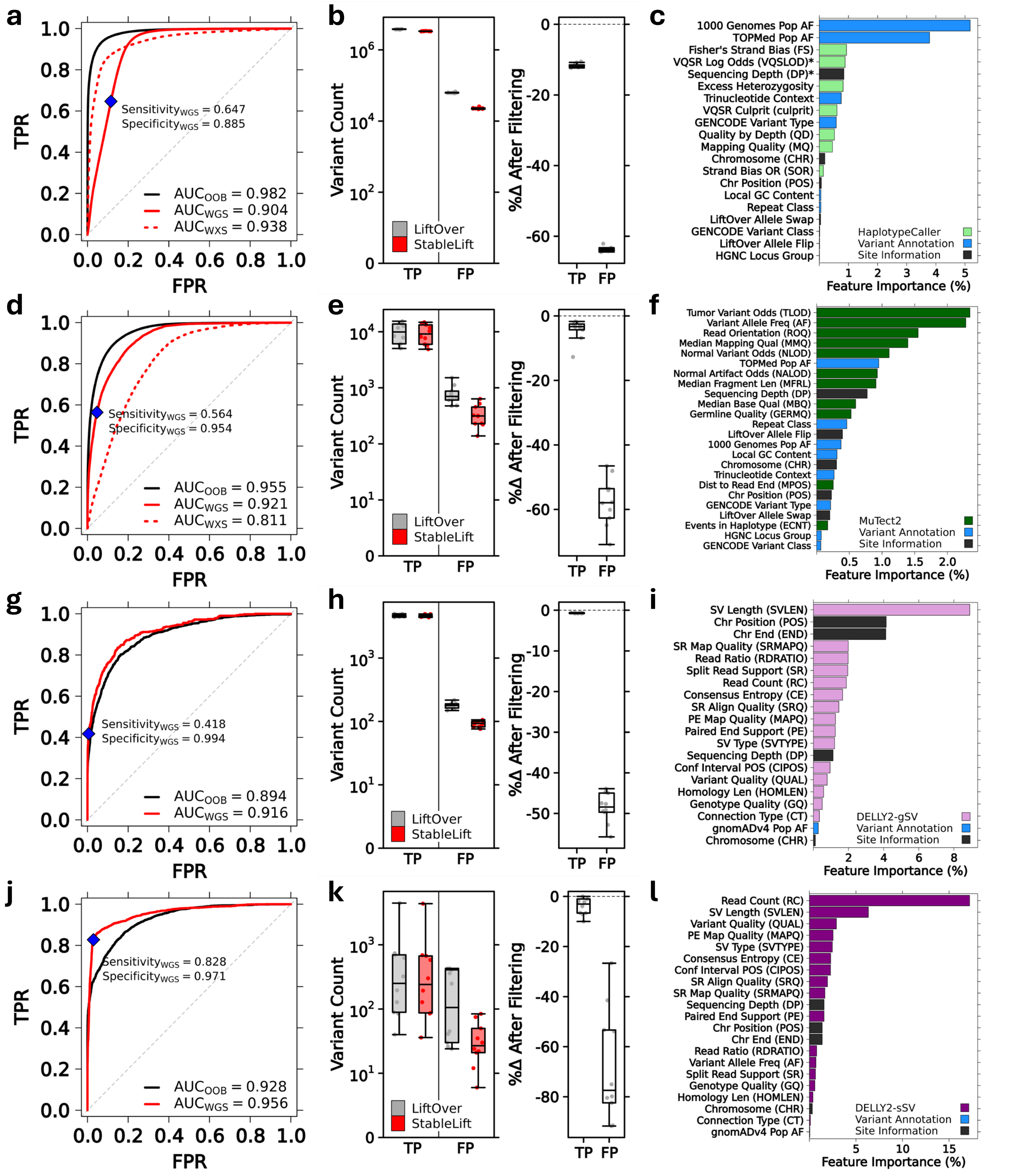


**Supplementary Figure 12.** GRCh38→GRCh37 StableLift model metrics. **a)** Random forest model performance for gSNPs (HaplotypeCaller) shown as ROC curves and AUC measures for out-of-bag whole genome training (OOB, solid black), whole genome validation (WGS, solid red), and whole exome validation (WXS, dashed red) sets. Default operating point maximizing F1-score highlighted (blue) with corresponding sensitivity and specificity in the whole genome validation set. **b)** Comparison of concordant (TP) and discordant (FP) gSNP counts before and after default StableLift filtering. **c)** Random forest feature importance colored by caller-specific metrics, variant annotations, and site information. **d-f)** Same as a-c for sSNVs (MuTect2). **g-i)** Same as a-c for gSVs (DELLY2). **j-l)** Same as a-c for sSVs (DELLY2).

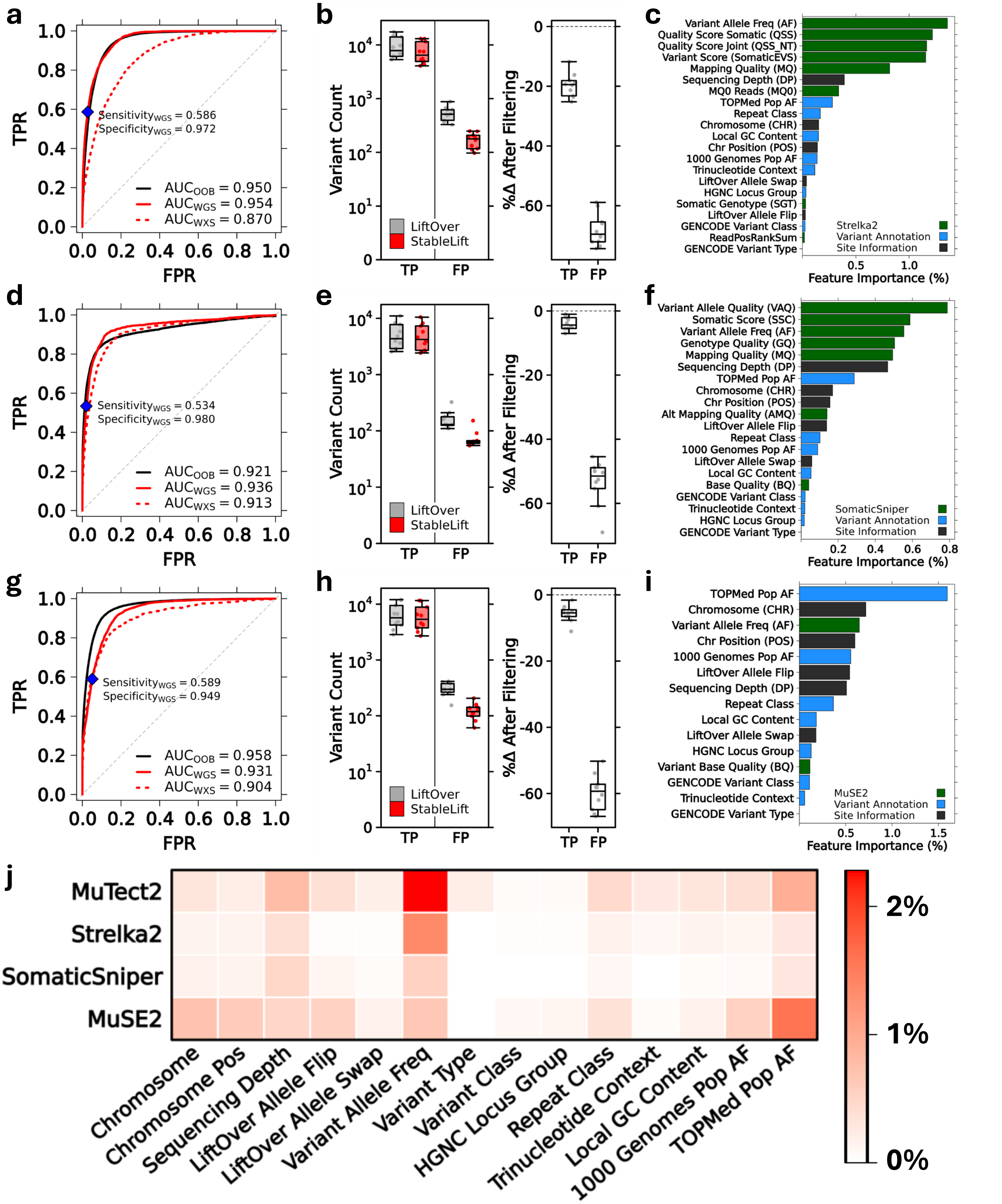

**Supplementary Figure 8.** GRCh38→GRCh37 StableLift model metrics. **a)** Random forest model performance for Strelka2 sSNVs shown as ROC curves and AUC measures for out-of-bag whole genome training (OOB, solid black), whole genome validation (WGS, solid red), and whole exome validation (WXS, dashed red) sets. Default operating point maximizing F1-score highlighted (blue) with corresponding sensitivity and specificity in the whole genome validation set. **b)** Comparison of concordant (TP) and discordant (FP) sSNV counts before and after default StableLift filtering. **c)** Random forest feature importance colored by caller-specific metrics, variant annotations, and site information. **d-f)** Same as a-c for SomaticSniper sSNVs. **g-i)** Same as a-c for MuSE2 sSNVs**. j)** Comparison of feature importance scores for features shared across the four sSNV callers.

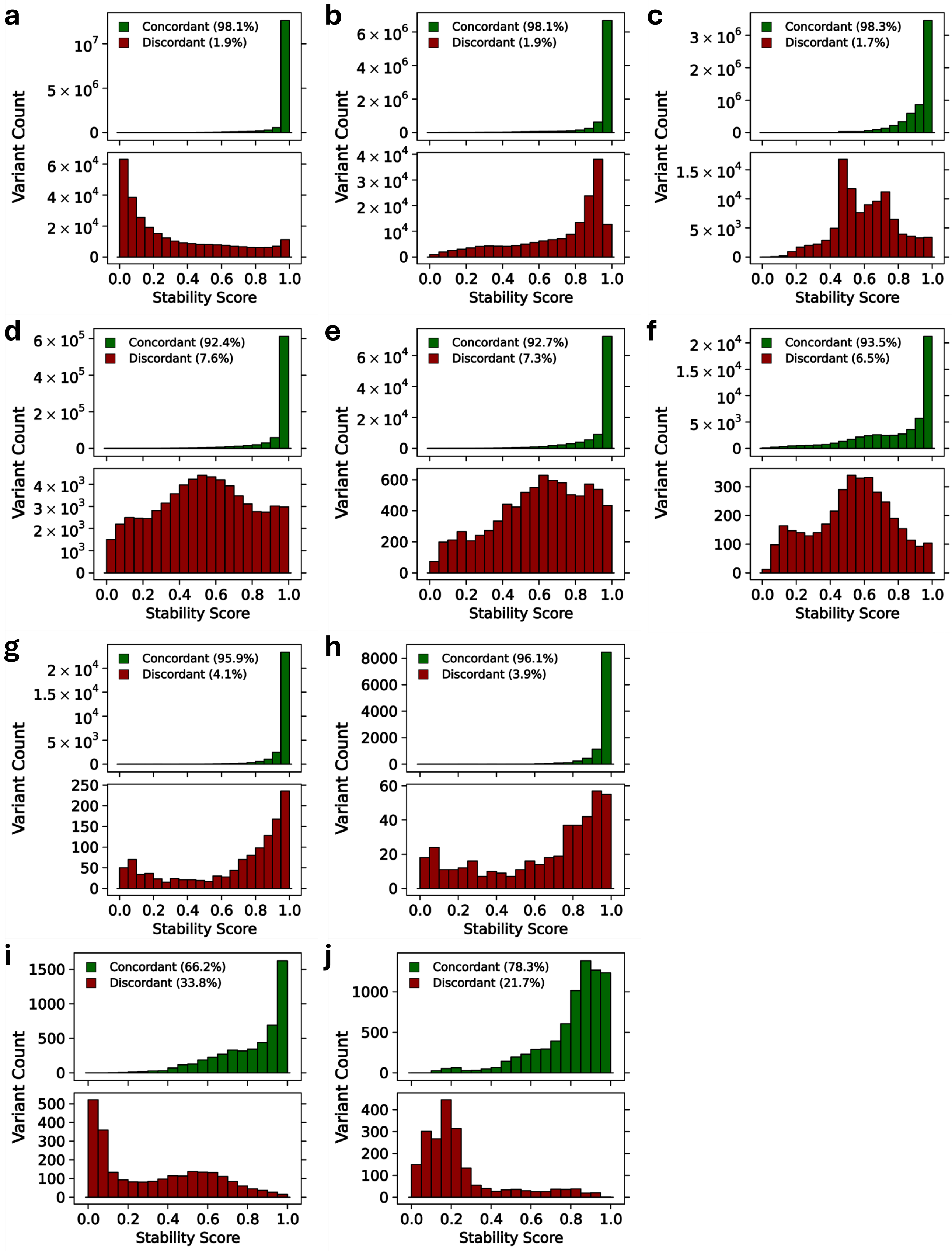


**Supplementary Figure 14.** GRCh38→GRCh37: Stability Score distribution of concordant *vs*. discordant variants (relative class proportions in parentheses) per variant caller. **a-c)** Concordant *vs*. discordant HaplotypeCaller gSNPs in whole genome training set (n=50), whole genome validation set (n=10), and exome validation set (n=60), respectively. **d-f)** Same as a-c for concordant *vs*. discordant MuTect2 sSNVs. **g-h)** Same as a-b for concordant *vs*. discordant DELLY2 gSVs. **i-j)** Same as a-b for concordant *vs*. discordant DELLY2 sSVs.

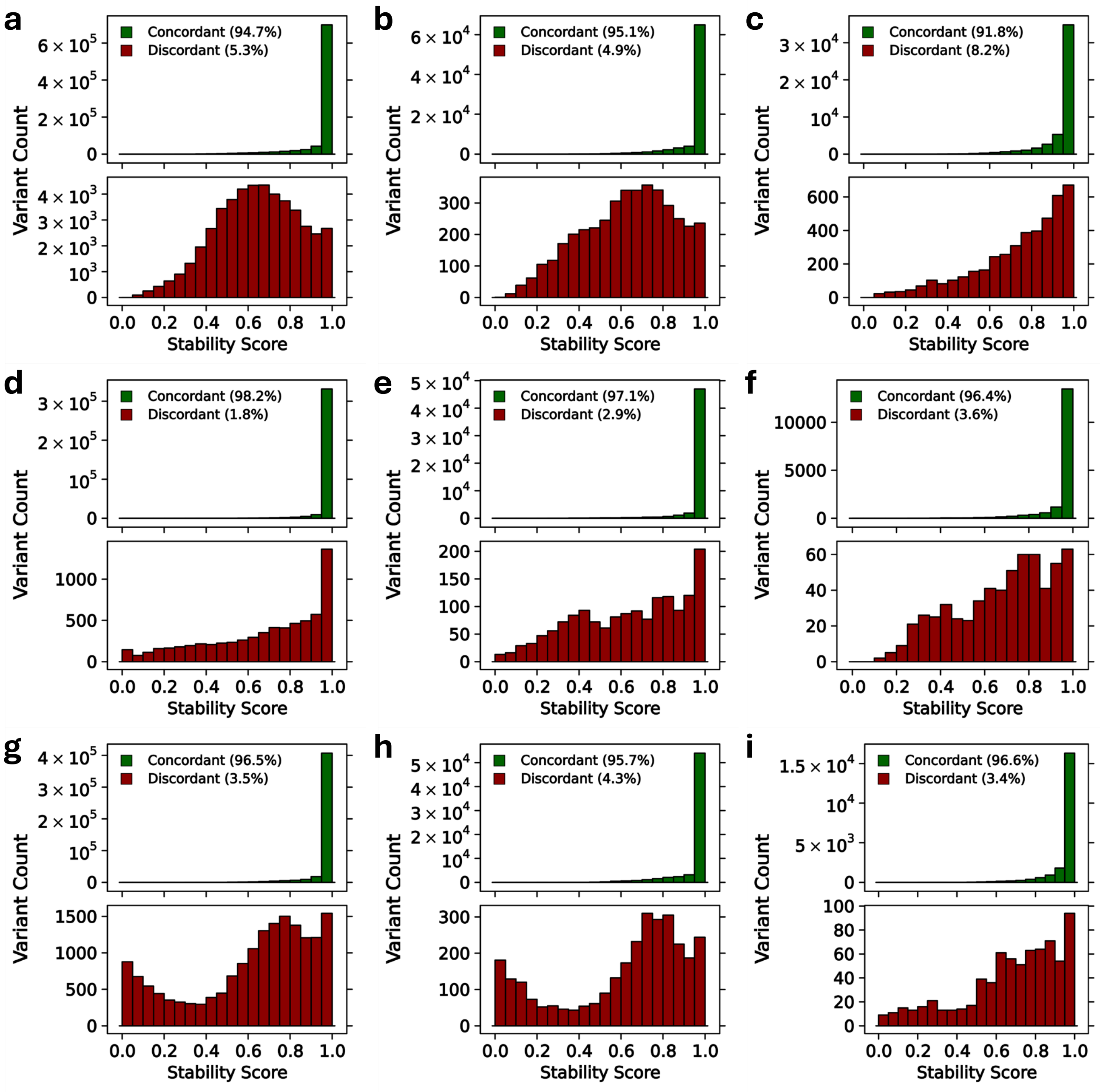


**Supplementary Figure 15.** GRCh38→GRCh37: Stability Score distribution of concordant *vs*. discordant variants (relative class proportions in parentheses) per variant caller. **a-c)** Concordant *vs*. discordant Strelka2 sSNVs in whole genome training set (n=50), whole genome validation set (n=10), and exome validation set (n=60), respectively. **d-f)** Same as a-c for concordant *vs*. discordant SomaticSniper sSNVs. **g-h)** Same as a-c for concordant *vs*. discordant MuSE2 sSNVs.

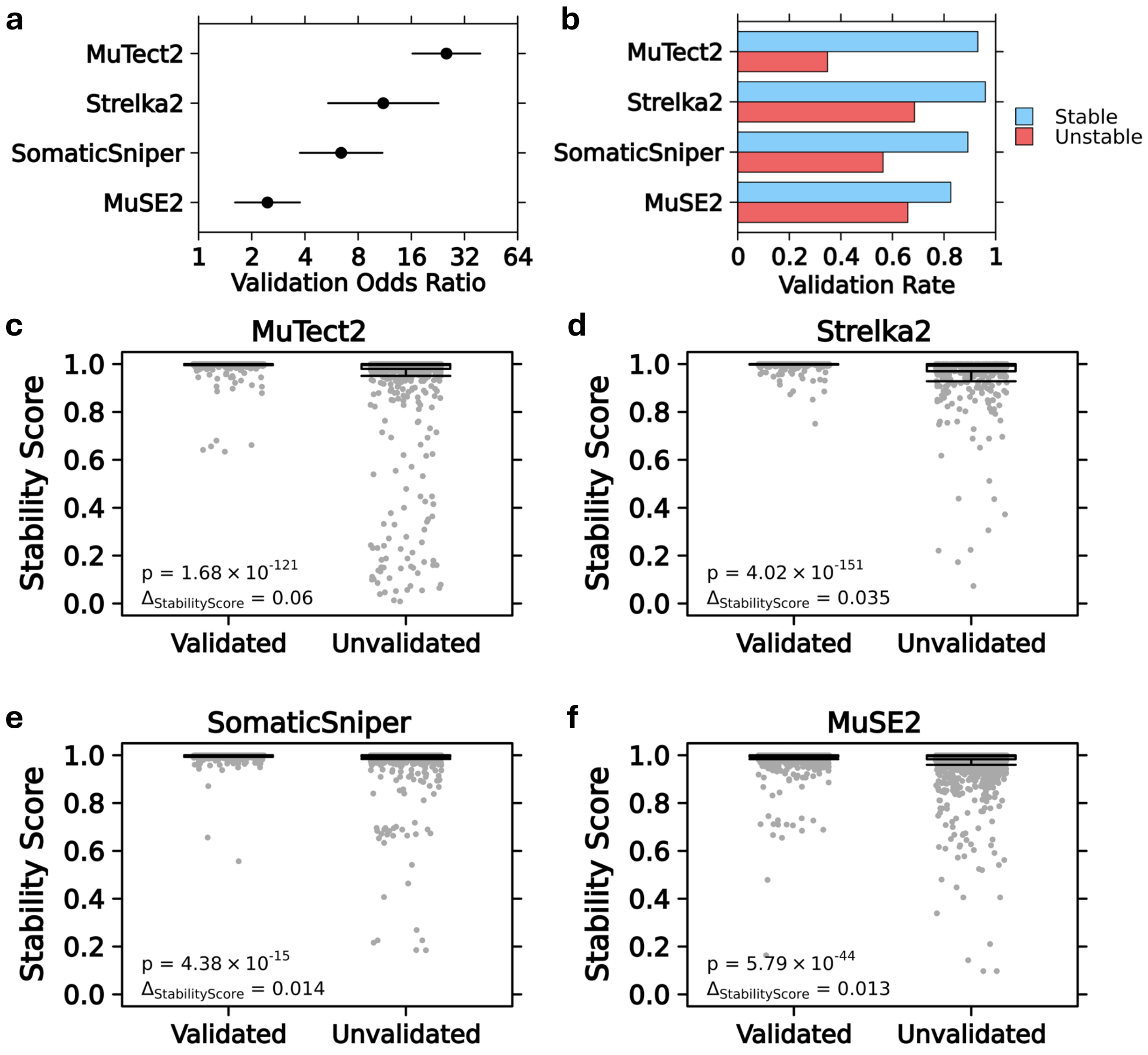


**Supplementary Figure 16.** GRCh38→GRCh37: Validation of StableLift on sSNV calls from five whole genome pairs with deep targeted resequencing. **a)** Validation odds ratio with 95% confidence interval of predicted "Stable" *vs*. "Unstable" variants for each sSNV caller. Validation status determined by concordance with variant calls from targeted deep-sequencing. **b)** Validation rate of predicted "Stable" *vs*. "Unstable" variants for each sSNV caller. **c-f)** Distribution of Stability Scores for validated *vs*. unvalidated sSNVs. Stability Scores per variant caller with Wilcoxon test p-value and difference in mean Stability Score shown. Strip plot downsampled to normalize group sizes for visualization.
